# Biregional division angles generate sharp apex and concave joints in leaves

**DOI:** 10.1101/2024.07.28.605531

**Authors:** Zining Wang, Yasuhiro Inoue, Atsushi Mochizuki, Hirokazu Tsukaya

## Abstract

Leaf apex, the distal end of the leaf blade, exhibits enormous shape variations across plant species. Among these variations, the sharp apex, characterized by its pointed tip, is important in species identification and environmental adaptation. Despite its taxonomic and ecological importance, the developmental mechanisms underlying the formation of a sharp apex remain unknown. The present study aims to investigate the curvature patterns and morphogenesis of the sharp apex to uncover these mechanisms using *Triadica sebifera* leaves. Our research revealed that the sharp apex marks the maximum positive curvature, and is flanked by concave joints with negative curvatures, suggestive of differential tissue growth and spatially regulated cellular behavior. Through a combination of wet experiments and numerical simulations, we demonstrated that biregional cell division angles, rather than locally differing cell expansion or division frequency, play a determining role in shaping distinct leaf morphology. Our study highlights the importance of spatiotemporal regulation of cell division angles during leaf development, suggesting that a biregional growth pattern and cellular behavior contribute to diversity in leaf apex morphology.

## Introduction

Leaves exhibit a wide range of shapes and structures, with their diversity being pivotal for photosynthesis and adaptation to local niches (*Nicotra et al., 2011*). Leaf morphology also plays an important role in taxonomy and systematics (*Viscosi et al., 2011*). To study the leaf morphology, it is essential to examine leaf contours, which could be described by a curve along the margin (*Tsukaya et al., 2018*). The quantification of leaf contour is usually achieved by curvature, which measures the degree to which a curve deviates from straight (*Liu et al., 2010*). In leaves, the contour can be divided into apex and base, which are often linked by joint regions (*Rickett, 1956*).

The leaf apex, which is situated at the distal end of the leaf blade, is crucial for classifying leaf shape (*Bong et al., 2014*). Classical plant anatomy categorizes apex shapes into three basic forms: pointed or sharp (acuminate, acute, cuspidate, and mucronate), smooth (obtuse and truncate), and concave (retuse, emarginate, and obocordate) (*Rickett, 1956*). In addition to their structural diversity, which has been utilized for species identification, leaf apices also perform essential physiological functions such as light signal detection, innate defense, and rain drainage (*Küpers et al., 2023*; *Juliana et al., 2015*; *Wang et al., 2020*). While the growth of the smooth apex in *Arabidopsis thaliana* can be explained by the common leaf development model (*Fox et al., 2018*), the mechanism of sharp-tipped apex morphogenesis, as shown in the leaves in Fig. 1, remains unknown. This knowledge gap raises fundamental questions regarding the developmental mechanisms that drive the growth of sharp apices in leaves.

**Figure 1.**
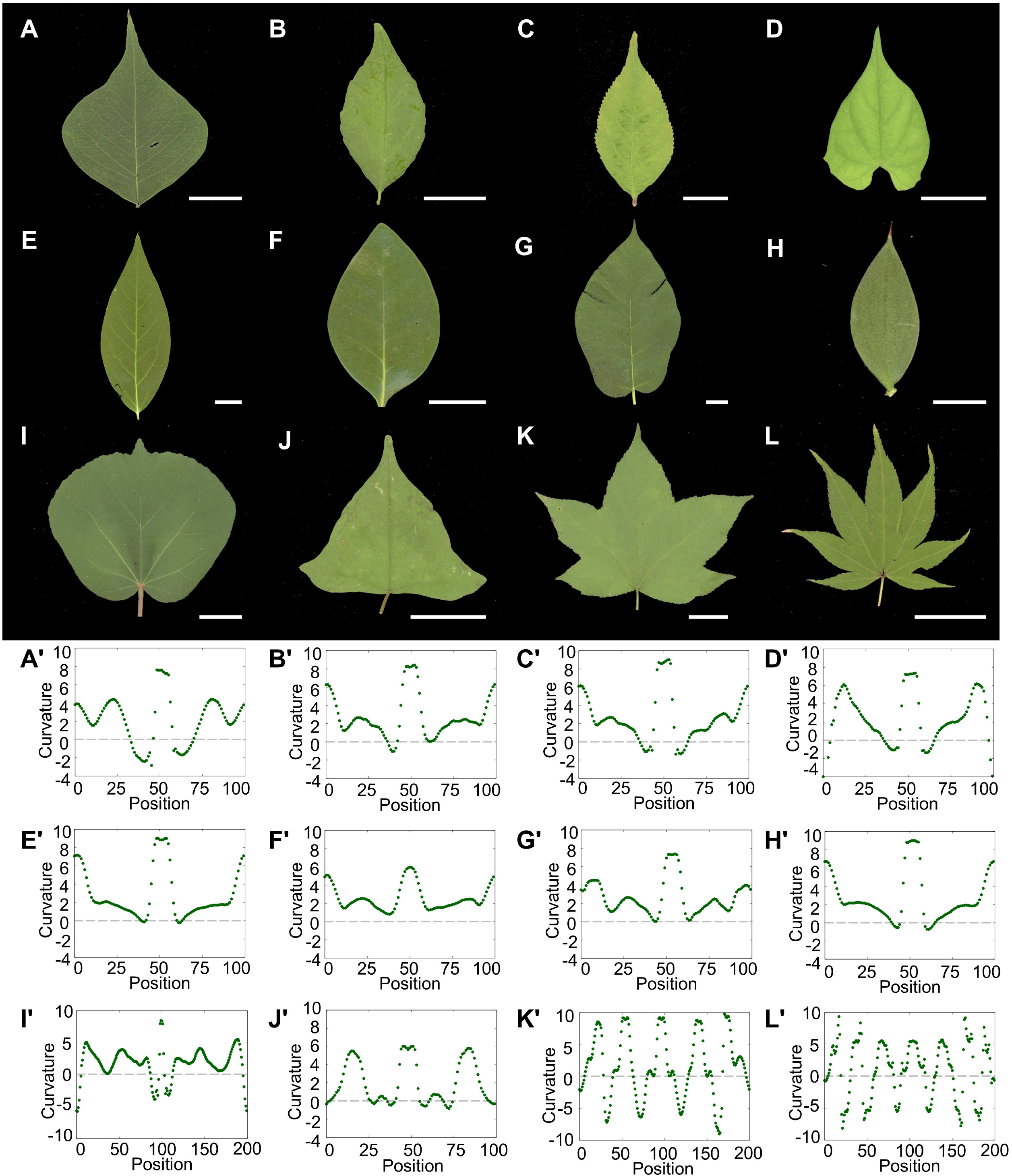
Leaves with characteristic apices and their curvature patterns. Leaves with acuminate apex: (A) *Triadica sebifera* L, (B) *Lagerstroemia indica* L., (C) *Callicarpa japonica* Thunb., and (D) *Ipomoea coccinea* L. Leaves with acute apex: (E) *Chimonanthus praecox* (L.) Link and (F) *Distylium racemosum* Siebold & Zucc. Leaves with cuspidate/mucronate apex: (G) *Ficus erecta* Thunb, (H) *Ruscus aculeatus* L., and (I) *Hibiscus hamabo* Siebold & Zucc. Leaves with multiple tips: (J) *Persicaria delibis* (Meisn.), H.Gross ex W.T.Lee, (K) *Liquidamabar styraciflua* L., and (L) *Acer palmatum* Thunb. (A’ -L’) Corresponding curvature patterns of (A-L) leaves. The curvature calculation starts from the base of the leaf and follows clockwise to enclose the contour. All leaves showed the peak of maximum curvature in the center, which corresponds to the apex. The peaks are always flanked by two troughs of minimum curvature, corresponding to the joint regions. Scale bars: 2 cm (A-G, I-L); 0.25 cm (H).

To examine the sharp apex morphogenesis, *Triadica sebifera*, which possesses an ideally sharp apex, was chosen as the research material in the present study. *T. sebifera*, also known as the Chinese tallow tree, is an important member of the spurge family (Euphorbiaceae). Its oil production, leaf color changes, ecological adaptation, and genome sequence has been well studied (*Liu et al., 2024*; *Carrillo et al., 2012*; *Luo et al., 2022*). Despite its important economic value, research about its leaf development has been absent to date. The leaves of *T. sebifera* exhibit an acuminate apex, rounded base, and concave joint region (Fig. 1A). These characteristics in shape make it an ideal model for apex growth and concave morphogenesis research. Because its rounded base could be easily explained by homogeneous growth (*Kennaway and Coen, 2019*), our study could focus on the morphogenesis of its sharp apex.

The common approach to study plant organ morphogenesis is to illustrate the detailed stages of organ development, map cell division and expansion patterns, and identify key genetic regulators (*Tsukaya, 2013*; *Du et al., 2018*; *Fox et al., 2018*). And due to the complexity of organ development, computational analysis and simulation have been widely applied to advance the understanding, as demonstrated by many previous studies (*Cheng et al., 2023*; *Hernandez-Lagana et al., 2021*; *Goh et al., 2023*). Our previous research of Arabidopsis petals and sepals also exemplified such methodology (*Kinoshita et al., 2022*). In this previous study, we first identified two stages of petal and sepal development, where in the early stage, cell division occurred uniformly across the entire primordia, and during the later stage, the region of active cell division shifted apically in petals and basally in sepals. By developmental experiment and computational simulation, we proved that such changes of cell division regions are critical for the differentiated morphogenesis into obovate and ovate shapes.

In this study, to understand the morphogenesis of apical growth and concave formation, we examined the roles of cell expansion and division during different developmental stages in the leaf primordia of *T. sebifera*. We propose a tissue-level biregional growth hypothesis and then complemented it with three cellular-level hypotheses: biregional cell expansion, biregional cell division angle, and biregional cell division frequency. Both wet experiments and 2D-vertex model simulations were used to test these hypotheses. Then, further generalizations were made from this case study of *T. sebifera*, with the aim of understanding the general morphogenesis process that generates various leaf apices.

## Results

### 1. Sharp apices exhibit the maximum curvature and are usually flanked by concave joints

Mature leaves with sharp apices were collected for shape and curvature analyses (Fig. 1 and S1). According to *Rickett (1956)*, sharp apices can be classified into three types according to their size and curvature pattern: 1. acuminate, sharply pointed, “implied a change in the direction of curvature from convex to concave” (Fig. 1A-1D, 1J-1L); 2. acute: “terminating gradually in a sharp point” (Fig. 1E, 1F); 3. mucronate or cuspidate: “tipped with a short abrupt point, when the midrib is produced beyond the apex in the form of a small point” (Fig. 1G-1I). Curvature analysis showed that all sharp apices were marked by a maximum positive curvature peak. These peaks were typically flanked by concave joint regions, which correspond to the negative curvature (Fig. 1A’-1I’). We also examined the curvature pattern of leaves with multiple tips (Fig. 1J-1L, 1J’-1L’). In the case of Japanese maple (*Acer palmatum*) (Fig. 1L, 1L’), seven lobes align with the seven curvature peaks and the indented edges between lobes matched the curvature local minimums. These pronounced changes in curvature suggest significantly differential growth in tissues and spatially regulated cell behavior. In addition, some leaves may exhibit a rather smooth curvature pattern and are not concave in the joint region, as shown in the case of *Distylium racemosum* (Fig. 1F, 1F’). These differences in the curvature patterns of apices and joints underscore the diversity of leaf apex morphogenesis across species.

To elucidate the mechanisms behind common curvature formations in leaves, *T. sebifera* was selected for its characteristic shape: acuminate apex, concave joint, and rounded base (Fig. 1A, 1A’). Curvature analysis of *T. sebifera* mature leaves revealed a common curvature pattern shared by sharp-tipped leaves. Specifically, in the basal region, the curvature measured approximately 2, indicative of its rounded shape; the joint region displayed a negative curvature, reaching a minimum of -2.67; and the apical region reached a maximum curvature of close to 8, highlighting the sharpness of the apex (Fig. 1A’). The circular basal part can be easily explained by homogeneous growth (*Lecuit and Le Goff, 2007*), enabling us to focus on the sharp apex and concave joint morphogenesis. This pattern of shape and curvature makes *T. sebifera* an ideal model for studying the general rules of apex growth and concave formation in subsequent experiments and analyses.

### 2. Differential tissue growth in the apex and base drives the formation of sharp apices and concave joints in leaves

We assumed that a sudden change in curvature could be generated by a difference in growth between the apical and basal regions and that the transition region of the curvature pattern could be the boundary of such differential growth. Based on this assumption, a tissue-level biregional growth model was made for the leaf morphogenesis in *T. sebifera* and sharp-tipped leaves: (1) the sharp apex could be generated by vertical growth (Fig. 2A, 2A’); (2) the circular base could be generated by isotropic growth (Fig. 2B, 2B’); and (3) the concave joint region could be generated by the difference in growth pattern between vertical growth in apex and isotropic growth in base (Fig. 2C, 2C’). With such growth patterns, we assumed that the sharp apex and concave joints of *T. sebifera* leaf could be generated (Fig. 2D, 2D’).

**Figure 2.**
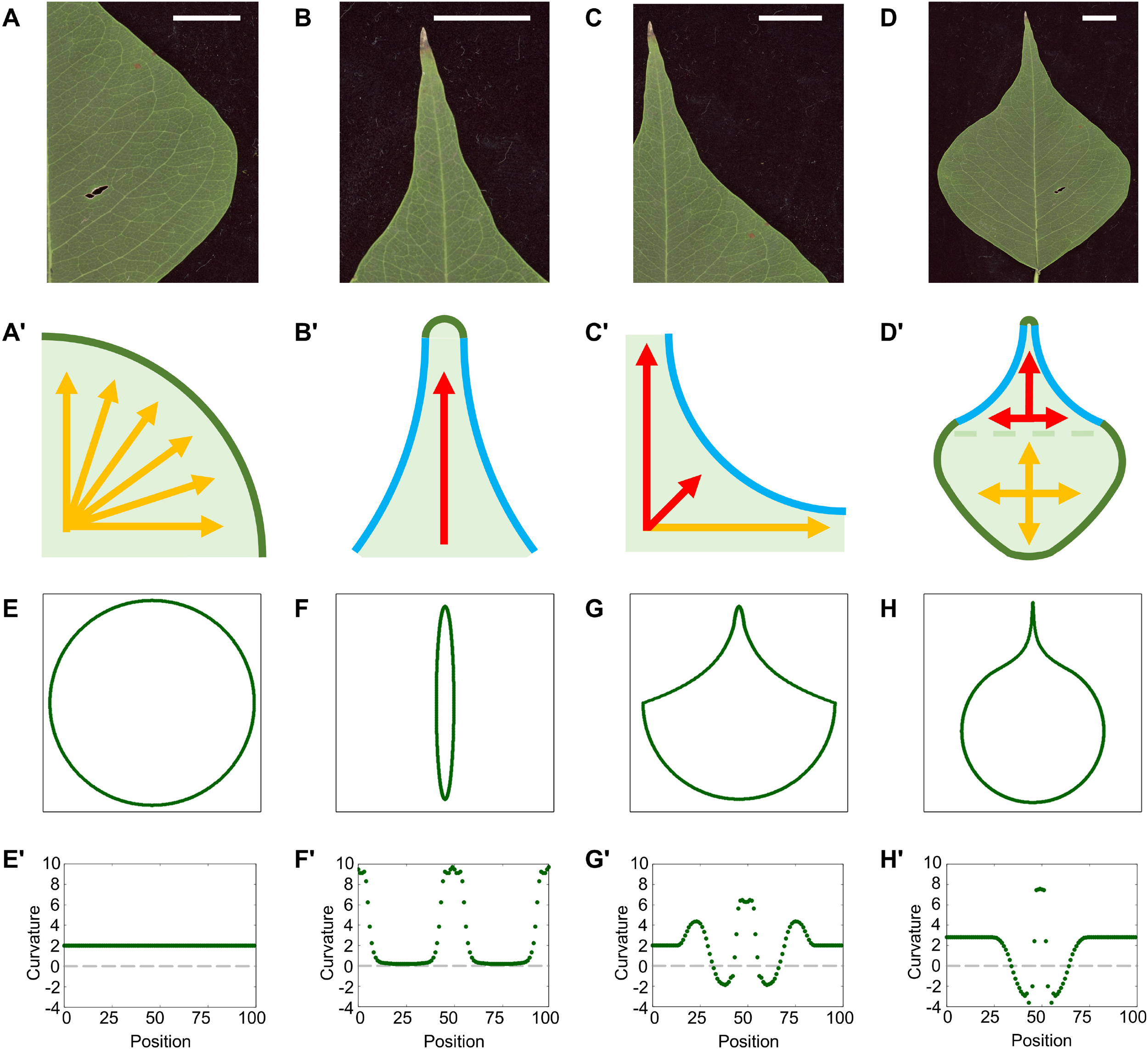
Tissue-level hypothesis of *Triadica sebifera* sharp-tipped leaf morphogenesis. (A, A**’**) Rounded basal region generated by homogeneous growth of leaf tissue. (B, B**’**) Sharp-tipped apical region generated via vertical growth of leaf tissue. (C, C**’**) Concave joint region generated via the differential growth between the vertically growing apex and homogeneously growing base. (D, D**’**) Desired organ shape in *T. sebifera* generated via biregionally differential growth. (E-H) simulation results of contour growth mapping generated via homogeneous growth setting, vertical growth setting (β = 0.8), biregional growth setting (β_*x*_ = 0.60, β_*y*_ = 1.00, *y*_*a*_ = 0.90 *y*_*b*_ = 0.5), and fine-tunning biregional growth setting (β_*x*_ = 0.8, β_*y*_ = 1.10, *y*_*a*_ = 1.00, *y*_*b*_ = 0.90). (E’-H’) Corresponding curvature pattern of (E-H). Scale bar 1 cm.

A simple growth model, named as contour growth mapping, was designed to test whether such hypotheses could generate the desired organ shape. This mapping abstracts the organ into a group of contour points and defines the growth of organs using a growing matrix that transforms the contour points. Our contour growth mapping with isotropic growth setting generated a circular organ (Fig. 2E, Movie S1) and vertical growth setting generated an elliptical organ (Fig. 2F, Movie S2). Between the vertically growing apex and homogenously growing base, a negative curvature formed in the joint region (Fig. 2G, Movie S3). Furthermore, by fine-tuning such growth patterns, contour growth mapping generated similar organ shapes and curvature patterns in *T. sebifera*, supporting the tissue-level biregional growth hypothesis (Fig. 2H, Movie S4).

This tissue-level argument is a rough phenomenological description that lacks cell-level details. To understand the mechanism of this sharp apex formation, we focused on the cellular behaviors that could realize tissue-level growth. We propose three cellular-level hypotheses based on biregional growth: (1) cell expansion is biased towards the proximo-distal axis in the apical region and cell expansion is less biased in the basal region (biregional cell expansion hypothesis, Fig. 3A), (2) cell division angles are biased in the apical region but random in the basal region (biregional division angle hypothesis) and (3) cell division frequencies are lower in the apical region and higher in the basal region (biregional division frequency hypothesis). Although the tissue-level hypothesis is straightforward, cellular-level hypotheses must be examined carefully through both experiments and simulations.

**Figure 3.**
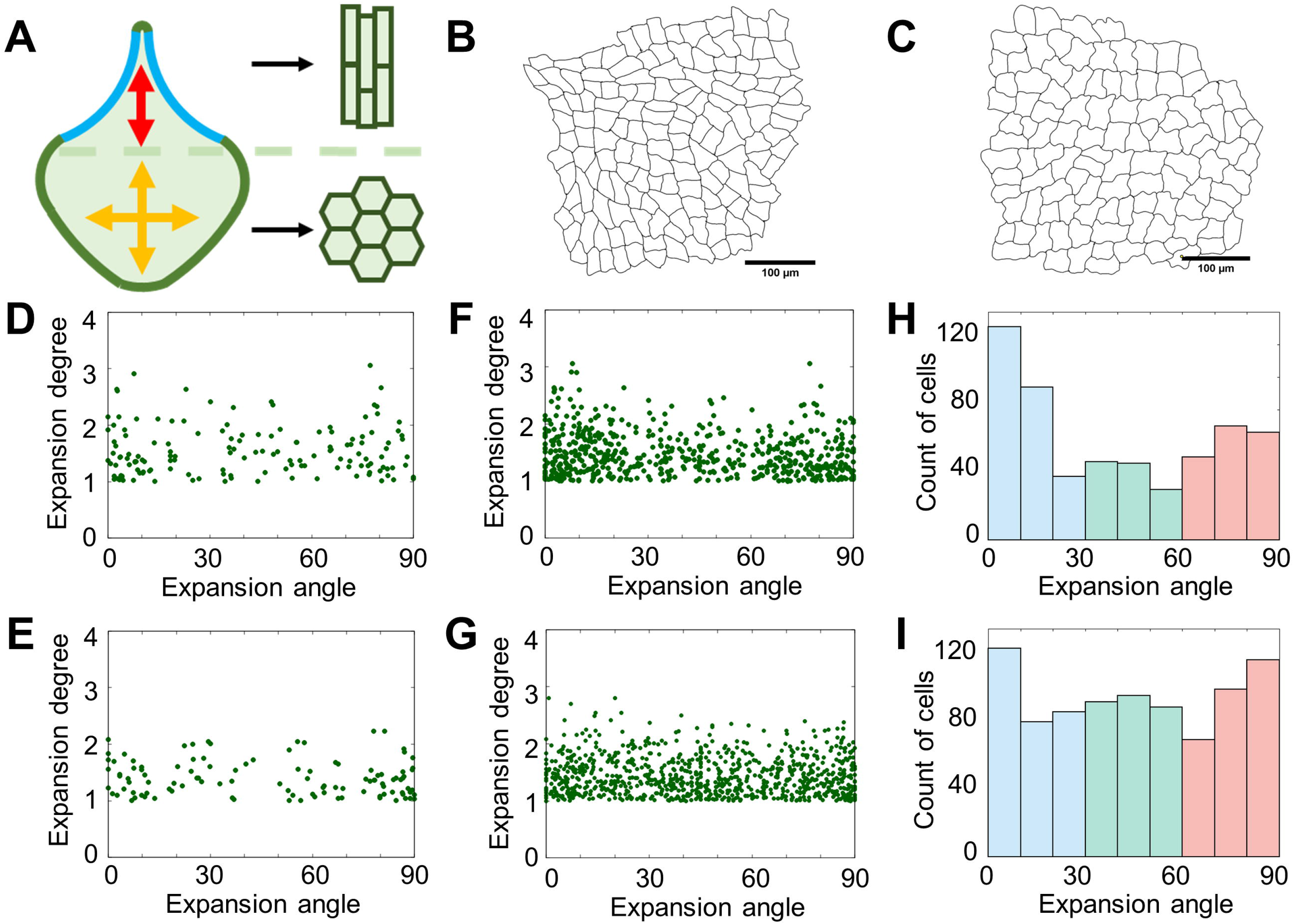
Biregional cell expansion hypothesis and its examination. (A) Diagram of biregional cell expansion hypothesis, where cells in apical region are biasedly expanded in proximal-distal axis and cells in basal region are less biasedly expanded. (B, C) Examples of adaxial epidermal cell morphology in apical and basal regions of mature leaves. (D, E) Distribution of biased cell expansion in (B) and (C), respectively. Distribution of biased cell expansion: (F) adaxial epidermal cells in the apical region, (G) adaxial epidermal cells in the basal region, (H) histogram of adaxial epidermal cells expansion in the apical region, and (I) histogram of adaxial epidermal cells expansion in the basal region. Cell numbers: (B, D) 145 cells; (C, E) 109 cells; (F, H) 591 cells; and (G, I) 903 cells.

### 3. No biregional cell expansion was observed in *T. sebifera* leaves

We first examined the biregional cell expansion hypothesis by observing cell shapes in *T. sebifera* leaves (Fig. 3B, 3C). This hypothesis suggests that cells in the apical region show polarized expansion along the proximodistal axis and cells in the basal region show less biased expansion (Fig. 3A). To observe whether such biased cell expansion occurred during *T. sebifera* leaf development, both mature and young leaf samples were fixed and cell walls were stained using Calcofluor White (mature leaf length: 5–7 cm; young leaf length: approximately 1 cm). The stained samples were then observed under a confocal microscope, and the contours of the epidermal and palisade cells were extracted (Fig. S2A-S2D). The cell contour was fitted to the smallest rectangle. The expansion of the cells can be represented by the shapes of these fitted rectangles, which are given by the angles of the major axes of the fitted rectangles, and the degrees of expansion are given by the ratios of length to width in the fitted rectangles.

Using the above methods, we analyzed the cell shapes in the apex and base of young leaves, but could not observe the predicted cell expansion pattern (Fig. S3). Even the apex of the leaf showed no polarized cell expansion along the proximodistal axis (Fig S3A, S3B, S3D). Similarly, in mature leaves, the cells in the apical region were aligned along the central-lateral axis (Fig. 3D-3I). Further detailed analyses of different regions of the leaf confirmed that there was no expected polarized cell expansion in the apical region (Fig. S2).

Thus, the observed biased expansion in both the apical and basal regions could not be the cause of sharp apex formation. The cell profiles shown in Fig. S3A suggested that division is a key factor contributing to sharp apex morphogenesis and no polarized cell shape was observed. In summary, the biregional biased cell expansion hypothesis was rejected based on the real cell morphology from a wet experimental perspective.

### 4. Biregional cell division angles were observed in *T. sebifera* primordia

The second hypothesis was tested using the EdU pulse-chase method to detect cell division angles (*Yin and Tsukaya, 2016*) in *T. sebifera* primordia. EdU pulse-chase experiments for different sizes of *T. sebifera* primordia were conducted. According to the EdU results, the primordia development could be divided into three stages: stage I, smaller than 1,000 µm in length (Fig. 4A, 4B); stage II, 1,000 to 2,500 µm (Fig. 4C); and stage III, larger than 2,500 µm (Fig. 4D). The division angles for stage I were biased in the vertical direction in both the basal and apical regions (Fig. 4E, 4F). In stage II, the cells divided in the vertical direction in the apical region and a random direction in the very basal region (Fig. 4G, 4H). For stage III, the EdU signal could only be observed in the basal part (Fig. 4D), which suggested that the arrest front appeared and cell division gradually stopped in the leaves (*Kazama et al., 2010*). These biased and random angle distributions were confirmed using the chi-squared test. Thus, a spatiotemporal pattern of biased cell division angles was observed in leaves, as expected from the biregional cell division angle hypothesis. However, it remains unknown whether such division angles can generate a sharp apex and a concave joint.

**Figure 4.**
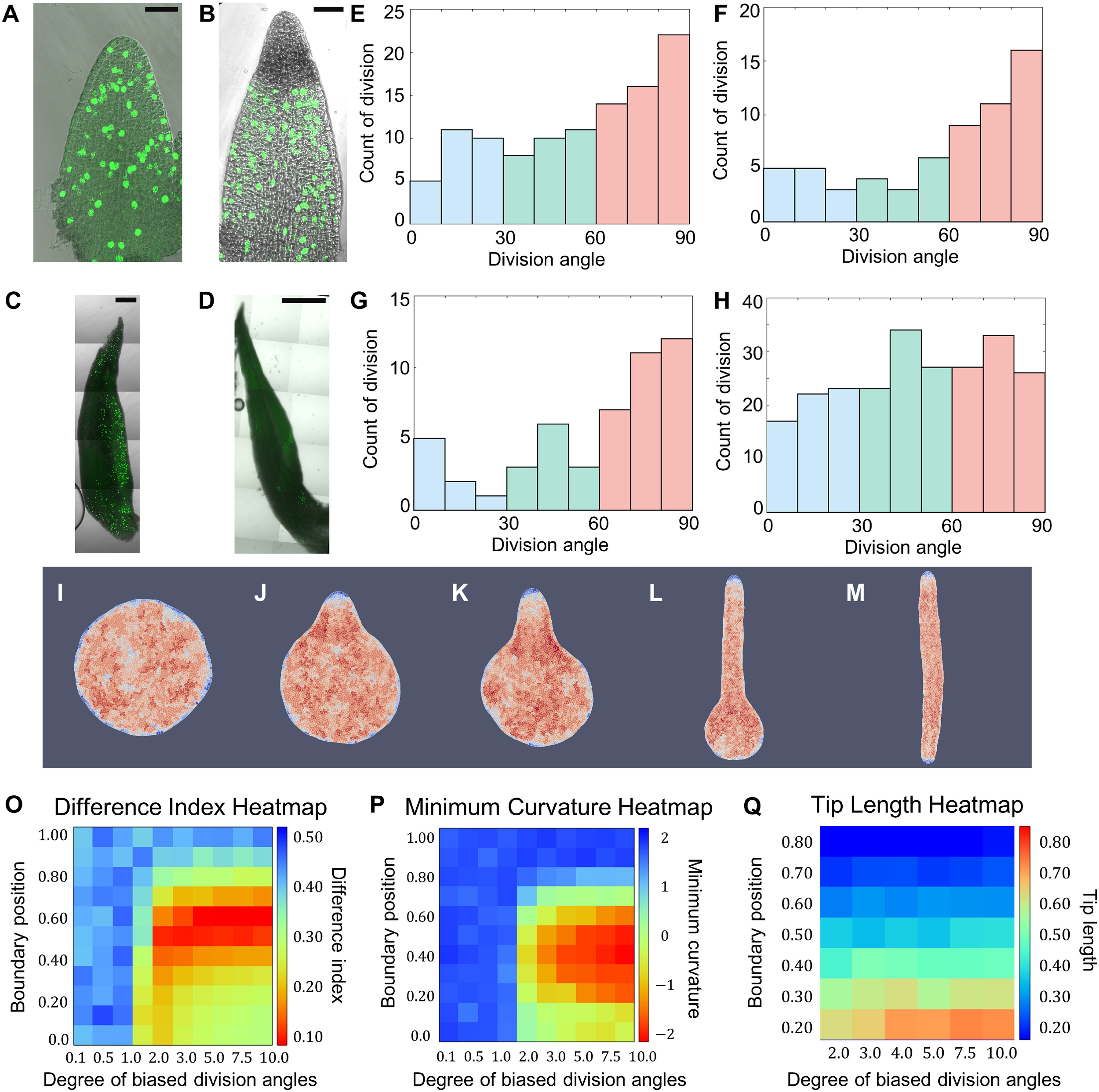
Examination of biregional division angles hypothesis. Leaf primordia stained by EdU pulse-chase method and observed using a confocal microscope: (A) Stage I, primordia length 348 µm; (B) Stage I, primordia length 490 µm; (C) Stage II, primordia length 2089 µm; and (D) Stage III, primordia length 2834 µm. Only the very basal part of the leaf has division signals. (E) Distribution of cell division angles in the apical region of leaf primordia in stage I (107 cell division events from 8 primordia). (F) Distribution of cell division angles in the basal region of leaf primordia in stage I (106 cell division events from 8 primordia). (G) Distribution of cell division angles in the apical region of leaf primordia in stage II (50 cell division events from 7 primordia). (H) Distribution of cell division angles in the basal region of leaf primordia in stage II (232 division events from 7 primordia). (I-M) Examples of organs generated from numerical simulations. The color indicates division counts: red indicates higher and blue indicates lower division counts. The corresponding boundary position and degree of biased division angles in the apical region are 0.50, 0.10 (I); 0.70, 10.0 (J); 0.60, 10.0 (K); 0.30, 10.0 (L); 0.00, 10.0 (M). (O) Heatmap of minimum curvature on *y*_*b*_, 1/*σ* space. (P) Heatmap of difference index on *y*_*b*_, 1/*σ* space. (Q) Heatmap of tip length on *y*_*b*_, 1/*σ* space. Scale bars: (A, B) 50 µm; (C) 200 µm; and (D) 500 µm.

### 5. Biregional cell division angles could generate sharp apex and concave joints in leaf

Next, we examined whether the observed biased cell division angles could generate sharp apex and concave joints in *T. sebifera* leaves. To construct such a causal relationship, we adopted vertex model simulation (*Nagai and Honda, 2001*), which can simulate organ growth from the given behavior of cells. In a previous study, we successfully used a vertex model simulation to generate obovate and ovate shapes of Arabidopsis petals and sepals based on their apical and basal meristem positions, respectively (*Kinoshita et al., 2022*). The model framework is the same as that used in a previous study. The biased cell division angles were implemented in the simulation by assuming that the division angles of cells followed the Gaussian distribution 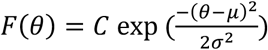on the angle space (0° < *θ* < 180°), where *C* is a normalizing constant. The direction and degree of cell division angle bias was given by the parameter *μ* and 1/*σ*, respectively in this Gaussian function.

In stage II primordia, the basal cells divided in a random direction and apical cells in the vertical direction. We fixed *μ* to be 90^°^ (vertical direction) in the apical region and examined the parameters 1/*σ* and *y*_*b*_, where *y*_*b*_ defines the boundary position that divides the apical and basal regions. An exhaustive parameter search was conducted on the 1/*σ, y*_*b*_ parameter space and a variety of apex shapes were generated (Fig. 4I-4M, Fig. S4). The extreme cases include the circular shape, where 1/*σ* ≈ 0 or *y*_*b*_ = 1, so no biased angles were exhibited (Fig. 4I, Movie S5, located at the middle-left part of heatmaps), and the rod shape, where *y*_*b*_ = 0 and 1/*σ* ≫ 0, so only vertical cell division occurred (Fig. 4M, Movie S6, located at the lower-right part of heatmaps). Apices of different lengths and sharpness emerged between the parameter ranges of these two extreme cases (Fig. 4J-4L, Movie S7-9, Fig. S4). The sharpness of the apex was mainly controlled by the degree of biased division angles (1/*σ*). The length of the sharp apex, given 1/*σ* ≫ 0, was mainly controlled by boundary between apex and base (*y*_*b*_).

To examine whether our simulation results were similar to the real organ in terms of contour, we used the “difference index,” which quantifies the difference between organ shapes (*Kinoshita et al., 2022*). The difference index analysis of the mature leaves of *T. sebifera* suggested that different leaves are not completely identical in shape, although they share the same characteristic curvature patterns (Fig. S5A, S5B). The threshold for defining such differences was set as 0.10 for difference index and an averaged contour was created for comparison with the simulated organs (Fig. S5C-E). The difference index analysis of our batch simulations suggested that)1/*σ, y*_*b*_) = (10,0.6) has the highest similarity with real leaves of *T. sebifera* (Fig. 4K). The results near this parameter choice passed the difference index threshold (red region in the heat map of Fig. 4O). The Gaussian fitting of our experiment-measured biased cell division angles suggested that 1/*σ* ≈ 2.89 in real leaf, which corresponds to the result that also passed the difference index threshold. Furthermore, curvature analysis of the simulated organs showed a negative curvature in the same parameter region (Fig. 4P). The length of the apex seemed to be unaffected by the degree of biased division angles, given such bias was significant enough to make sharp apices (1/*σ* > 2.0). The apex length was determined using the boundary position between the vertically dividing apical region and the randomly dividing basal region (Fig. 4Q). In summary, our vertex model simulation results confirmed that the experimentally discovered biased division angles could be a causal factor in the morphogenesis of sharp apices and concave joints. Thus, the experimental evidence supported a prerequisite for the second hypothesis (biased division angles) and a causal relationship was established through simulation analysis. The temporal pattern of the biased cell division angles is examined in a later section.

### 6. Biregional cell division frequency could not generate the desired leaf shape

As discussed by *Tsukaya (2018)*, a lower cell division frequency in the very early stage of leaf primordia, where the apical part is made, might contribute to sharp apex formation. Based on this assumption, we examined the theoretical possibility that a biregional biased cell division frequency could contribute to the sharp apex and concave joint formation using a vertex model simulation. Two parameters were used: *y*_*b*_, the position of the boundary that divides the apical and basal regions, and *f*_*r*_, the relative division frequency of the apical and basal regions. The exhaustive parameter search and analysis did not reveal the desired organ shape and curvature pattern using the defined settings (Fig. 5A-5E, S6). The difference index and minimum curvature analyses did not reveal shapes that passed the success threshold (Fig. 5F, 5G). Our previous study on Arabidopsis petals and sepals (*Kinoshita et al., 2022*) suggested that a differential cell division frequency controlled by a Gaussian distribution cannot generate a sharp-tipped shape. This frequency control can create anisotropic shapes, such as obovate and ovate (Fig. 5B, 5E, Movie 10S), but not sharp-tipped shapes, which are more irregular and might require more differential growth. In summary, the third hypothesis was rejected by the simulations.

**Figure 5.**
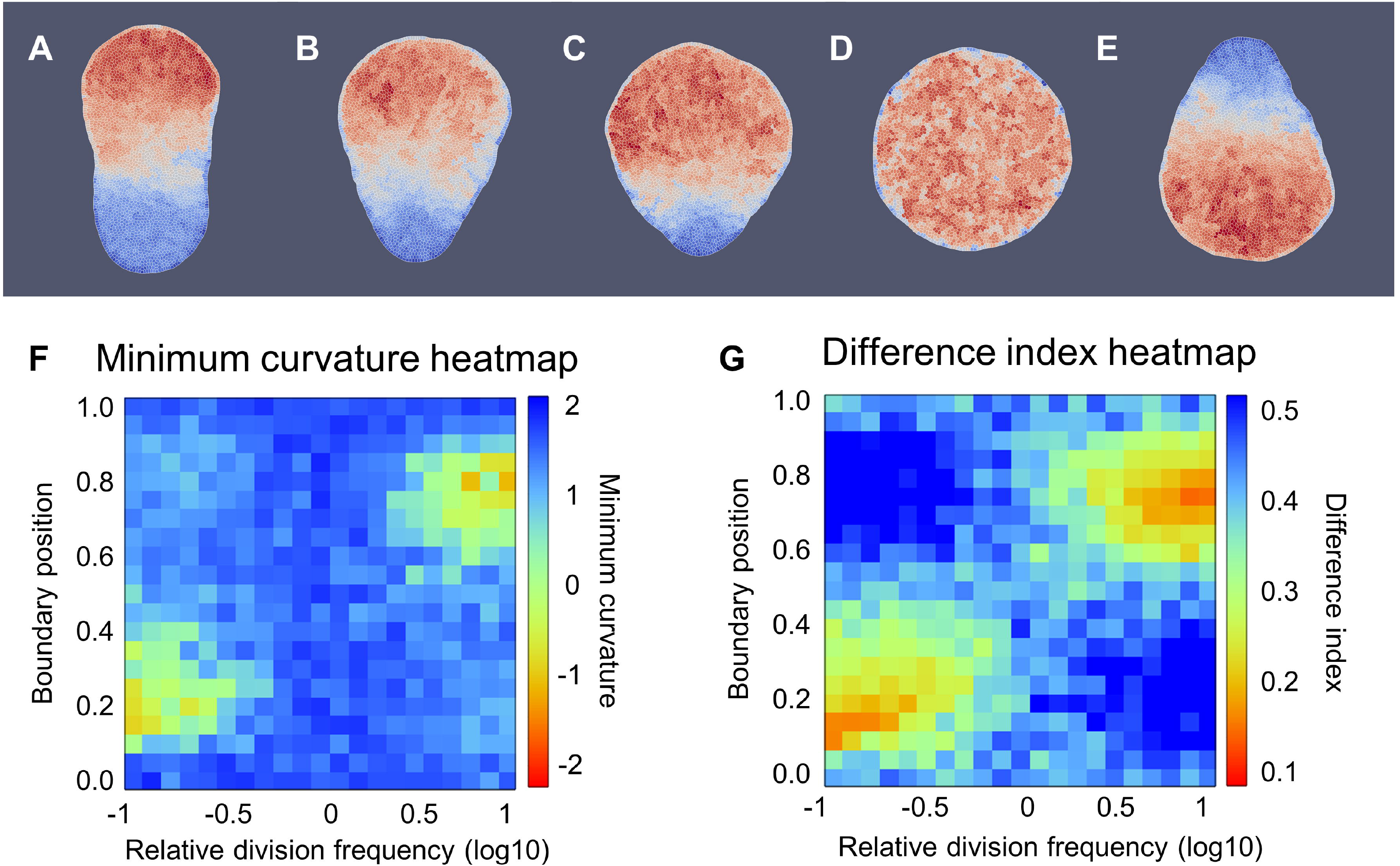
Analysis of biregionally biased division frequency simulation results. Characteristic results of biregional division frequency simulation: (A) *y*_*b*_ = 0.8, *rf* = 0.13, oval shape; (B) *y*_*b*_ = 0. 7, *rf* = 0.1, obovate shape; (C) *y*_*b*_ = 0.6, *rf* = 0.1, shape transitioning from obovate to rounded; and (D) *y*_*b*_ = 0.5, *rf* = 1.0, rounded shape; (E) *y* = 0.3, *rf* = 10.0, ovate shape. (F) Heatmap of minimum curvature on *y*_*bou*nd_, *rf* parameter space. (G) Heatmap of difference index on *y*_*bou*nd_, *rf* parameter space. No simulation results from this setting could pass the success threshold (difference index<0.10, minimum curvature<-2).

### 7. Gradual boundary of biregional division angles generates smooth organ shape

The biregional division angle hypothesis was suggested by both experiments and simulations. We further investigated this hypothesis by considering the gradual boundaries and temporal patterns of the biased division angles. In our previous simulations, we defined the boundary as a sudden change in the cell division angle pattern. Here, we examined whether a continuous change in the differential cell division angles could generate the desired organ shape. The organs were then divided into apical, intermediate, and basal regions. These three regions were divided by introducing two parameters, *y*_*basal*_ and *y*_*apical*_. The bias angle increased linearly in the intermediate region. Our simulation results suggested that more gradual boundaries generated smoother organ shapes and less negative curvatures (Fig. 6 and S7). The results obtained by gradual boundary control may correspond to other leaves with a smooth concave joint, as shown in Fig. 1S. But in the case of *T. sebifera*, which possesses a sharp concave joint, a rather abrupt boundary might be required.

**Figure 6.**
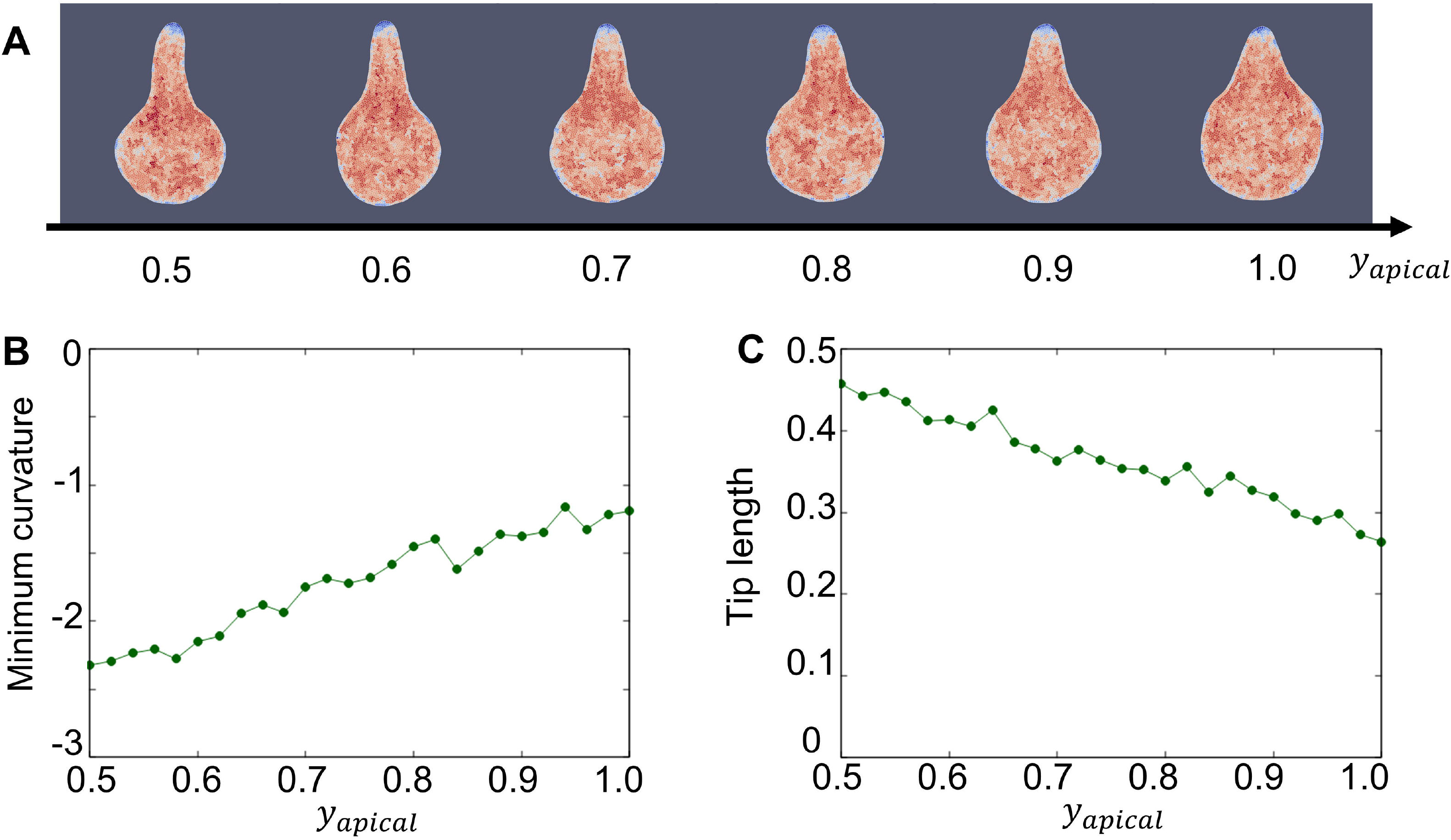
Simulation results and analysis of biregional division angles setting with gradual boundary. (A) Simulation results of growth boundary growth with *y*_*basal*_ fixed to 0.5. (B) Analysis result of minimum curvature of (A). (C) Analysis result of tip length of (A)

### 8. Temporal controls of biased cell division angles could also generate the desired sharp apex

As shown in Fig. 4A-4D, in stage I of *T. sebifera* development, the cells divided across the primordia and were biased in the vertical direction. The primordial shape appears to be elliptical. In stage II, the cell division angles emerged as a biregional pattern. In stage III, the cells ceased to divide from the apex to the base. However, these temporal patterns require further investigation. A vertex model using a temporal setting was designed to test the morphogenetic effects of these cell division phenomena. In this setting, as demonstrated in Fig. 4B, we intentionally used only the temporal control of cell division angles and combined it with the restriction of the cell division region in the later stage by the arrest front.

The two division patterns (arrested front and temporally biased cell division angles) were examined separately. The simulation results for the arrest front alone generated a transition of organ shapes from circular to elliptical (Fig. 7A, 7C, Movie S11) and the timing of the transition from circular to elliptical was determined by the square of the size of the arrest front in our simulation (Fig. 7D). This suggests that the transition occurs at the time point when the cell number/(arrest front size)^2 exceeds some threshold value and that the size of the arrest front only affects the transition time but not the potential shape pattern. The simulation results for the temporal division angles alone generated organ shapes ranging from thin to thick ellipses (Movie S12). These data suggest that, if applied separately, the arrest front and temporally biased cell division angles cannot generate a sharp apex. However, the combined control of the arrest front and biased division angles generated the desired organ shape (Fig. 7B, 7E, Movie S13). The cell division angles averaged by the temporal axis showed that such temporal patterns had the same distribution as our previous spatial regulation and EdU pulse-chase experimental results (Fig. 7F, 7G). Further exhaustive parameter searches of the combined patterns generated successful results (Fig. S8). Thus, these simulation results suggest that spatial and temporal regulation have similar cell division angle distributions and both generate a sharp apex. The reality of *T. sebifera* and leaf morphogenesis in other species should be a spatiotemporal pattern involving a mixture of temporal and spatial regulation of cell division angles.

**Figure 7.**
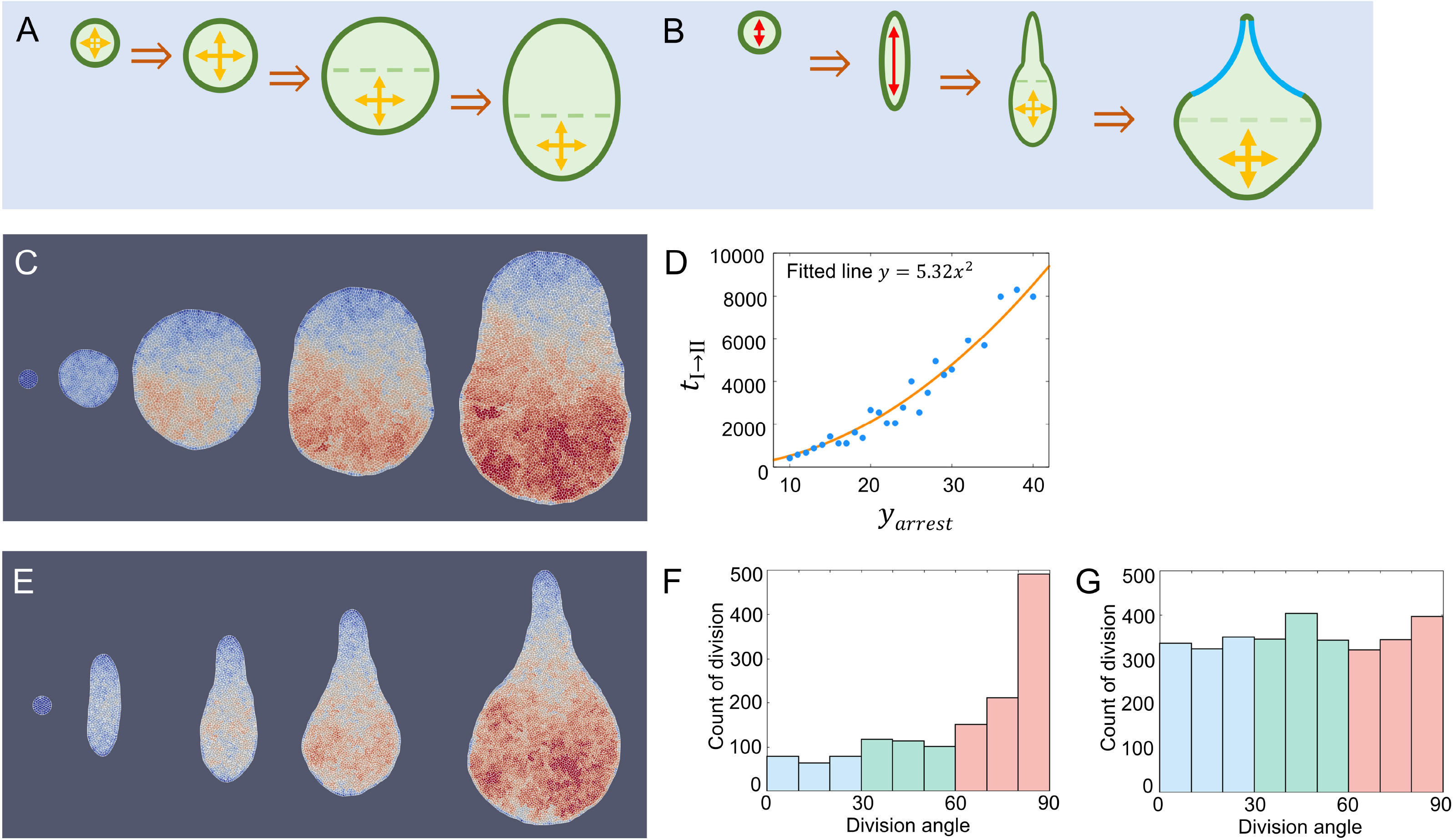
Temporal regulation of division angles. (A) Diagram of arrest front simulation. Initially, all cells divide in equal division frequency, later only basal cells can divide. The division angles are random. (B) Diagram of simulation setting combined arrest front and temporal division angles. Initially, all cells divide in the proximal-distal axis (vertical direction). Later, only basal cells can divide and the division direction is random. (C) A time series of arrest front simulation result example (*y*_*arrest*_ = 20) by setting of (A). In the early stage (stage I), the initial circular organ grew into a larger circle. In later stage (stage II), the circular organ grew into a cylindrical organ. The first two results were from stage I; the third was in the transition; and the fourth and fifth were in stage II. (D) Relationship between *y*_*arrest*_ and transition time from stage I to stage II can be fitted by *y* = 5.32*x*^2^, R^2^ = 0.955, which suggested that *y*_*arrest*_ can only affect the transition time from stage I to stage II but not the shape pattern itself. (E) Time series of successful simulations combined arrest front and temporal division angles, by setting of (B). The first two were corresponding to the initial elongation stage; The third was in the transition stage; The fourth and fifth were in the final stage of circular base growth. Division angle distribution averaged in time of (E): (F) apical region and (G) basal region. The division angle distribution showed a similar pattern to that of biregion division angle results and experiment results (Fig. 4G, 4H, and 4K).

## Discussions

### 1. Biregional division angles generate sharp apex

We examined the developmental mechanisms of sharp apices and concave joint formation. First, we quantified and classified the sharp apex and concave joints using curvature analysis. The curvature patterns of 28 leaves showed that a sharp apex corresponded to the maximum curvature flanked by concave joints (Fig. S1). Based on this shape and curvature difference between the apex and base, we proposed a tissue-level biregional growth model in which apical tissue followed vertical growth pattern and basal tissue had homogeneous growth pattern. Contour growth mapping based on this biregional growth setting successfully generated the desired organ shape, confirming the tissue-level biregional growth hypothesis (Fig. 2). The biregional growth hypothesis at the tissue level was subsequently subdivided into three cellular-level hypotheses: (1) biregional cell expansion hypothesis; (2) biregional cell division angle hypothesis, and (3) biregional cell division frequency hypothesis. For the biregional expansion hypothesis, we checked the cell morphology in *T. sebifera* but did not observe the predicted vertically biased cell shape (Fig. 3). Thus, the biregional expansion hypothesis was rejected. For the biregional division angle hypothesis, we measured the division angles using EdU pulse-chase and found that vertically biased division angles existed. We further established a causal relationship between the discovered division angle patterns and sharp apex formation using a numerical simulation (Fig. 4). For the biregional division frequency hypothesis, we found that differential frequency could only generate obovate or ovate shapes (Fig. 5, 6S), in accordance with our previous research on Arabidopsis petal and sepal morphogenesis (*Kinoshita et al., 2022*). Thus, only the biregional division angle hypothesis was validated by quantitative examination. The generalization of this research in terms of the intermediate region between the apex and base was examined, and we found that a more gradual boundary generated a smoother apex and a less concave joint (Fig. 6). In addition, a temporal pattern of the division angles abstracted from the experimental results could generate shapes similar to the spatial pattern (Fig. 7). By integrating both experimental and simulation results, we demonstrated that sharp-tipped leaves are produced through biregional division angles, likely achieved by temporal regulation rather than direct spatial regulation. The following sections discuss several aspects of these findings.

### 2. Why division angles, rather than expansion or division frequency, are chosen to form a sharp apex

Cell expansion along the proximal–distal axis, or cell elongation, is the main cellular mechanism of root elongation (*Goh et al., 2023*), submerged leaves (*Koga et al., 2020*), and leaf petioles (*Polko et al., 2012*). We expected that a similar mechanism might elongate the apex and sharpen its shape; however, the actual cell morphology did not show similar cell expansion patterns, as observed in the case of *T. sebifera* (Fig. 3). Interestingly, the cell shapes showed some bias towards the central-lateral and proximo-distal axes but not in the diagonal direction, suggesting that certain cell profiles existed during development. We also confirmed no vertically biased cell expansion in the leaves of *Ipomoea coccinea* (Convolvulaceae, Solanales) (Fig. 1D) and *Persicaria debilis* (Polygonaceae, Caryophyllales) (Fig. 1J), which have distant phylogenetic relationships with *T. sebifera* (Euphorbiaceae, Malpighiales) (Fig S9). These results suggest that, in general, cell expansion is not a strategy for sharp apex formation. Furthermore, differential expansion may generate an imbalance in mechanics, cell density, or cell identity between the apex and base, which is different from the relatively homogeneous elongation pattern observed in the cells of roots or submerged leaves. To maintain similar cell identity at both the apex and base of mature leaves, regionally biased cell expansion may not be a good strategy for creating a sharp apex.

Next, we discuss a potential approach to form a sharp apex using the biregional division frequency. Differential division frequency patterns contribute to serration formation (*Bilsborough et al., 2011*) and petal/sepal formation in Arabidopsis (*Peng et al., 2022*). As examined in Arabidopsis (*Kamaza et al., 2010*), leaf cells cease to divide in the apical region in later developmental stages. Thus, a sharp apex should form earlier and the base widens later. However, our simulation results showed that the differential frequency along the proximodistal axis could only generate obovate and ovate shapes, not the desired patterns of a sharp apex, removing the possibility of it being the causal factor.

Division angle control, which was found to be the causal factor of such sharp apex formation in this study, is also a determining factor for root elongation (*Hong et al., 2018*), thickening growth of unifacial leaves in *Juncus* (*Yin and Tsukaya, 2019*), and trap formation in *Utricularia gibba* (*Whitewoods et al., 2020*). These studies, along with ours, indicate that division angle control is responsible for the elongation or protrusion of organs with more drastic isotropic growth than division frequency control. When such protruding parts are folded, tubes and other concave structures can form, as seen in the carnivorous pitcher leaves of *Sarracenia purpurea* (*Fukushima et al., 2015*), and also in the sharp apex in *Triadica sebifera* in this study.

Cell expansion, division frequency, and division angle are applied by plants to generate ideal shapes (*Dupuy et al., 2010*). Our mission was to determine which was responsible for the morphogenesis of the target shape pattern. And in this study, we found that division angle control with specific spatiotemporal pattern was the causal factor in sharp-tipped pattern morphogenesis.

### 3. Potential adaptiveness and molecular mechanism of sharp apex formation

The potential evolutionary significance of a sharp apex has been examined in previous studies. *Küpers et al. (2023)* focused on the ability of the apex to detect light signals and found that a lower light intensity may promote petiole growth to reach out for potential light. Thus, a sharp apex may serve as an antenna for gathering environmental information during plant competition for light. *Wang et al. (2020)* and *Liu et al. (2023)* examined another potential function of the sharp apex: increasing water drainage. Although their research demonstrated the higher water drainage efficiency of leaves with sharp apices during heavy rain, *T. sebifera* does not live in high-precipitation environments. Therefore, water drainage efficiency might not be an evolutionary driver of this sharp apex formation. Further studies are needed to demonstrate the adaptive meaning of sharp apex in general (*Tsukaya, 2018*). The evolutionary meaning of a wide base is rather clear – larger area for photosynthesis. Thus, a sharp apex serving as an environment signal detection antenna and a wide base serving as a photosynthesis factory may be a good choice for plant leaves. The joint region linking the sharp apex and wide base can be naturally concave as shown in this research, but its evolutionary meaning remains unclear. In addition to a leaf lamina with sharp apex and widened base, a petiole or internode that is sufficiently long to avoid self-shading is also advantageous (*Weijschedé et al., 2006*). Our study shows that to generate such leaves, a biregional cell division angle control is adopted by leaves. The molecular mechanisms may include *ANGUSTIFOLIA*3 (*AN3*) (*Horiguchi et al., 2005*; *Kawade et al., 2013*) and auxin-related pathways, which require further investigations using methods such as RNA-seq (*Kinoshita and Tsukaya, 2022*).

### 4. Diverse forms of leaf apex and their potential morphogenesis mechanisms

The diversity of the leaf apex extends beyond its sharp-tipped shape. The apex morphology can vary, ranging from obtuse, as observed in Arabidopsis, to concave, as observed in species such as *Oxalis* and *Liriodendron* (Fig. S10A-S10E). Curvature analysis revealed a negative curvature at the apex of these leaves, with varying degrees or sizes of concavity (Fig. S10A’-S10E’). This concave apex formation may arise from meristem separation into right and left regions, with inhibited cell division in the middle region. To illustrate this hypothesis, a simple contour growth mapping was employed with this setting (Fig. S10F-S10H and Movies S14-S16). By controlling the size and degree of meristem splitting, we generated a series of concave apices corresponding to *Oxalis* species (Fig. S10F’-S10H’). And further experiments on cell division patterns are needed in future research.

In addition to variations in apex morphology, leaves may develop several tips in addition to the apex position, as shown by *Acer palmatum* (Fig. 1L). These tips were recognized as leaf serrations and might have adopted developmental mechanisms other than the sharp apex morphogenesis observed in this study. For leaf serrations, the division frequency is enhanced in the local auxin maxima along the leaf margin, resulting in a relatively decreased division frequency between the maxima and contributing to serration outgrowth (*Kawamura et al., 2010*). In compound leaf formation, such differential growth begins at a very early stage of leaf primordium formation, leading to leaflet separation during later growth stages (*Luo et al., 2024*). Various shape changes, such as simple leaves, shallow serrations, deep serrations, compound leaves, and twigs, can be attributed to similar modifications in the control of cell division (*Bhatia et al., 2021*). Our simulation results indicated that pure division frequency control leads to only mildly isotropic growth, which is insufficient to generate multiple drastic acuminate tips in the maple leaf (Fig. 6S). We expect that a combined strategy of inhibited division frequency between tips and biased division angles within the tips may give rise to this multi-tipped shape.

## Materials and methods

### 1. Plant materials and growth

Mature leaves without specific explanations were collected from the Botanical Gardens, Graduate School of Science, University of Tokyo, Tokyo, during late autumn, 2023. *Persicaria debilis* samples were collected from the Nikko Botanical Garden, University of Tokyo, in late autumn of 2023. Leaves from four *Oxalis* species were collected from the Hongo campus of the University of Tokyo in April 2024. *Ipomoea coccinea* seeds were collected from a population that invaded Kamakura, Kanagawa, Japan, and mature leaves were collected after 50DAS under laboratory cultivation. *Triadica sebifera* seeds were collected from the Botanical Gardens of the University of Tokyo, Japan. For the pre-sowing treatment, *T. sebifera* seeds were submerged in water for one day at room temperature to soften, followed by the removal of their protective white wax coats. The seeds were then preserved at 4°C for at least one month of dormancy breaking. After the vernalization, they were sown and grown at 30°C under constant light conditions (∼40μmol m^−2^s^−1^), with a regular supply of water containing 1 g/L power Hyponex (Hyponex). *Triadica sebifera* saplings were grown in a greenhouse at the Hongo campus of the University of Tokyo.

### 2. Curvature analysis of leaf contours

Mature leaves were scanned using a GT-X830 (EPSON) at 600 dpi in the TIFF format. ImageJ software was used to remove dust particles and other undesired artifacts from the background. The contours of mature leaves were extracted using ImageJ. Curvature calculation was based on the method described by Driscoll et al. (2012). Leaf contours were parameterized using 100 boundary points equally distanced along the contour via a piecewise linear interpolation method. For leaves with complicated shapes or small tips, such as *Acer plamatum*, 200 boundary points were parameterized to enhance the sensitivity and fully capture the contour details.

The curvature of each parameterized boundary point was calculated using the Kåsa circle-fitting algorithm (*Kåsa, 1976*). Circle fitting was performed using each boundary point and its two boundary points that are ten boundary points neighboring away. The boundaries were normalized by their lengths. The absolute value of the curvature was defined as the reciprocal of the radius of the fitted circle. The symbol of curvature was designated as positive if the center of the fitted circle was located inside the leaf and negative if the center was located outside the leaf. The curvature was smoothed by averaging the two neighboring boundary points. Curvature plotting began from the bottom of the leaf in a clockwise direction and concluded at the starting point. The minor-scale curvature contributed by the leaf teeth was recognized as noise compared to the large-scale curvature made by the sharp apex and was thus removed through averaging in the algorithm. Curvature analysis of the simulated leaves was conducted in the same manner using a circle-fitting algorithm with 100 boundary points.

Leaves with a sharp apex were classified based on the description by *Rickett (1956)*. All sharp apices correspond to the maximum positive curvature. An acuminate apex was defined as a sharp apex with concave joints. An acute apex is defined as a sharp apex without a concave joint. Mucronates and cuspidates are defined as sharp but small apices. Based on the discussion by Rickett, mucronates and cuspidates are indistinguishable and in our curvature analysis, the difference between mucronates and cuspidates seemed inapplicable.

### 3. Contour growth mapping

Contour growth mapping was used to test tissue-level growth hypotheses. In this model, the shape of the plant organ is represented by a series of points *P*_*i*_(*x*_*i*_, *y*_*i*_), *i* = 1,2, …, *n* on the contour. Organ growth was abstracted by the movement of these points and further represented as a contour growth matrix:

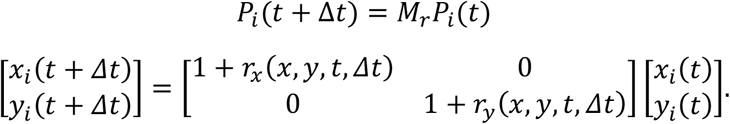

where the *P*_*i*_ (*x*_*i*_, *y*_*i*_) is the contour point *i* at time zero and 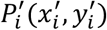 is the contour point *i* after time interval Δ*t. r*_*x*_ (*x, y, t*) defines the growth rate of the boundary point along the x-axis and *r*_*y*_ (*x, y, t*) defines the growth rate of *y*_*i*_ along the y-axis during this time interval.

For homogeneous growth, the contour growth matrix is given as

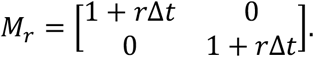

where *r* is a constant value, defining the growth speed of contour points.

For vertical growth, the contour growth matrix is given as

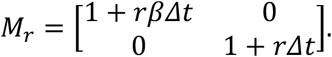

where *r* is the growth speed of contour points in the vertical direction and *β*(< 1) is the degree of biased growth speed of contour points in the horizontal direction.

For biregional growth with linear interpolation, the contour growth matrix is given as

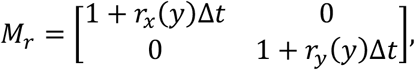

where *r*_*x*_ (*y*) is a piecewise function:

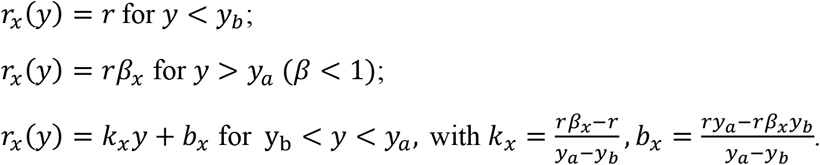

*r*_*y*_ (*y*) is a piecewise function:

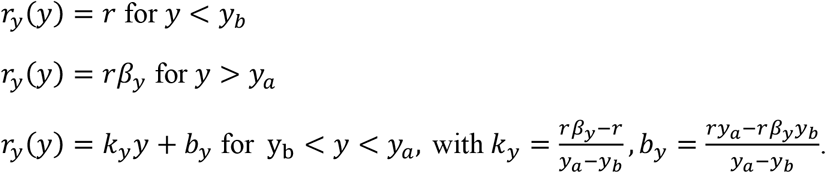

where *y*_*b*_ is the boundary of the basal region, *y*_*a*_ is the boundary of the apical region, *r* is the growth speed of the contour points in the vertical direction, and *β*_*x*_ and *β*_*y*_(< 1) are the degree of biased growth speed of contour points in the horizontal and vertical directions, respectively. This piecewise function implied homogeneous growth in the basal region, vertical growth in the apical region, and isotropic growth in the joint region.

For *Oxalis* species concave apex morphogenesis, the contour growth matrix is given as

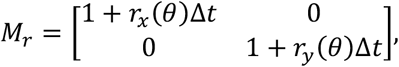

where *θ* is the angular parameter, indicating the angle of rotation from the positive x-axis in a polar coordinate system. We used this to determine the size of the inhibited growth region and the distance between the left and right meristems.

For 90° − *θ*_m_ > *θ*, this is the right meristem with a standard growth rate: *r*_*x*_ (*θ*) = *r*_*y*_ (*θ*) = *r*

For 90° + *θ*_m_< *θ*, this is the left meristem with standard growth rate: *r*_*x*_ (*θ*) = *r*_*y*_ (*θ*) = *r*

For 90° − *θ*_m_ < *θ* < 90° + *θ*_m_, this the inhibited growth region in the middle region. And for 90° − *θ*_m_ < *θ* < 90°, the growth factor is calculated by the sigmoid interpolation between (90° − *θ*_m_, *r*) and (90°, *rβ*):

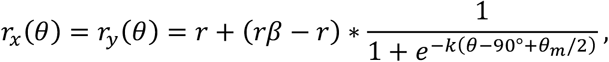

For 90° < *θ* < 90° + *θ*_m_, the growth factor is calculated by sigmoid interpolation between (90°, *rβ*) and (90° + *θ*_m_, *r*):

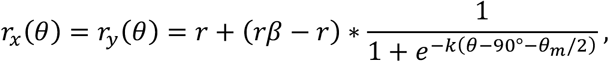

where *k* is the scaling factor that determines the steepness of the curve, *θ*_m_ is the distance between the left meristem and right meristem positions, and *β* is the degree of growth inhibition.

For all contour growth simulations, the initial contour was set as a circle with a radius of 5 and the contour was represented by 1,000 points, distributed evenly along the circumference. The simulation concluded in 100 steps. *r* was set as 0.0641, since (1 + 0.0641)^100^ ≈ 500, which is the expansion rate from early primordia (150 *μm*) to the mature leaf (7.5 cm), in terms of length. The other parameter values are indicated for each simulation. The simulation was conducted, analyzed using C++ codes, and plotted using Gnuplot.

### 4. Cell morphology observation and analysis

Mature leaves of *T. sebifera* were collected from the Koishikawa Botanical Garden in late autumn 2022. Pieces from the apical, joint, and basal regions were cut from mature leaves for observation. Young leaves were obtained from the seedlings. Cut pieces of mature or young leaves were washed with phosphate-buffered saline (PBS), fixed in FAA (37% formaldehyde, 5% acetic acid, 50% ethanol, all in volume/volume), and stored at 4°C. Subsequently, fixed samples were treated with 0.5 mg/L Calcofluor White (Sigma-Aldrich) diluted in ClearSee solution (*Kurihara et al., 2015)* for at least 2 days. The stained samples were mounted and observed under a confocal microscope FV3000 (Olympus).

The cell morphology from confocal microscope observation was manually extracted using ImageJ. Cells near veins and stomata were excluded because they exhibited special deformations (*Linh et al., 2018*). The cell area was calculated using ImageJ software, and biased cell expansion was analyzed using C++ code. Each cell contour was recognized as a polygon and fitted using a minimum rectangle. The length and width of the fitted minimum rectangle are defined as those of the cell. The degree of biased cell expansion was then calculated as the length-to-width ratio, while the angle of biased cell expansion was determined as the angle between the length and medial-lateral axes. The division angle along the proximodistal axis was defined as 90°.

### 5. Cell division angles detection and analysis

The protocol for the detection of cell division angles using EdU pulse-chase was modified from that of *Yin and Tsukaya (2021)*. Leaf primordia of *T. sebifera* were collected from seedlings and saplings. The samples were first washed with PBS for 30 min to remove wax. Subsequently, they were incubated in Murashige and Skoog inorganic salt medium (MS0; Wako) (*Murashige and Skoog, 1964*) supplemented with 10 µM EdU (Click-iT EdU Imaging Kit, Invitrogen) at 30°C under continuous light for 2 hours (pulse). The samples were then washed three times with MS0 for 5 min 3 times to remove excess EdU. The samples were then incubated under MS0 for 5 h 45 min at 30°C under continuous light (chase). Primordia were then fixed using FAA and stored at 4°C. Primordia were further dissected and detected using the Click-iT EdU Imaging Kit (Invitrogen) according to the manufacturer’s instructions. The samples were also stained with 0.5 mg/L Calcofluor White (Sigma-Aldrich) diluted in ClearSee for 1 d, washed with PBS three times for 5 min, mounted, and observed under a confocal microscope FV3000 (Olympus).

The criteria for choosing a valid pair of daughter cells were as follows: (1) the EdU staining patterns of the nuclei of the two daughter cells should be similar and (2) the shapes and sizes of the two daughter cells were similar. The center of the valid daughter cell nucleus was then extracted from the confocal images using ImageJ. Division angles were defined as the line connecting the two daughter cells and the medial-lateral axis of the leaf (90° along the proximo-distal axis). The chi-square goodness-of-fit test was performed in Python using the scipy.stats package. Gaussian fitting was used to quantify the biased cell division angle distribution using C++ code based on Guo’s algorithm (*Guo, 2011*).

### 6. Basics of vertex model simulation

To explore the cellular-level hypotheses of apex growth and concave formation, we employed a 2D vertex model to simulate organ morphogenesis, as initially outlined by *Nagai and Honda (2001)*, and further modified it in our previous work (*Kinoshita et al., 2022*). In this model, plant cells are abstracted as polygons and a plant organ is conceptualized as an assembly of these polygons that share edges with neighboring polygons. The deformation of the cells is represented by the movement of the vertices of these polygons, governed by a set of ordinary differential equations that determine the position of each vertex over time. Specifically, *r*_*i*_ represents the position vector of *i*-th vertex. The dynamic behavior of each vertex aims to minimize potential energy U, as defined by *Hayakawa et al. (2016)*.

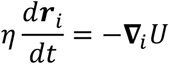

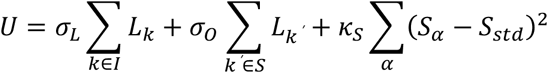

where *l*_*k*_ is the length of edge *k, l*_*k*′_ is the length of edge *k*^′^ at the tissue surface, *S*_α_ is the area of cell α, *S*_*std*_ is the standard cell area, sets *I* and *S* are the sets of vertices inside the organ and on the organ surface, respectively, *σ*_*L*_ and *σ*_*O*_ are the interface energy per unit length between two neighboring cells and at the tissue surface, respectively, and *K*_*S*_ is the elastic constant of the cell. Parameter values were set as follows: *σ*_*L*_ = 0.5, *σ*_*O*_= 2.0, *K*_*S*_ = 1.0, *S*_*std*_ = 3.5.

Cell division is modeled by introducing a new segment that bisects the cell area and adding two new vertices at the intersection points between the new segment and the boundary segments of the mother cell. This segment represents newly formed cell walls. The division angle is defined as perpendicular to this segment or the new cell wall and is either assigned randomly or determined by a predefined division angle control. The frequency of cell division was controlled using specific methods described later. This simulation began with an initial arrangement of 61 cells such that the entire mass of cells formed a circular boundary and progressed until a total of 3,000 cells. The results from the vertex model were visualized using the ParaView and Visualization Toolkit (VTK) library. Coding for the vertex model simulation and analysis was performed in C++, with line graphs and heatmaps plotted using Gnuplot, and videos produced with the OpenCV library.

### 7. Cell division angle and frequency control in spatial patterns

In our vertex model, the timing and orientation of cell division were precisely controlled to mimic biological processes. The potential of each cell to divide is tracked by a value called the cell time (*t*_*cell*_,_*i*_). During the initiation of the simulation, each cell was assigned a random number between 0 and a value called the cell-time threshold (*t*_*th*_). For every 10,000 numerical simulation steps, the cell time of each cell increases by 1, and for every 100,000 numerical simulation steps, cells with a cell time above the threshold (*t*_*cell*_,_*i*_ > *t*_*th*_) have a 1/4 chance of dividing. To control the division frequency carefully, two key modifiers were introduced for *t*_*th*_: the cell area modifier and a spatial modifier. The area modifier prevents the division of small cells and favours the division of larger cells, which is a common approach used in vertex modeling. For the biased cell division frequency hypothesis, the spatial modifier is defined as

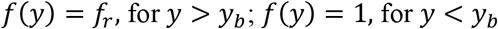

where y is the relative position of a cell along the y-axis, *y*_*b*_ is the boundary between the apical and basal regions, and *f*_*r*_ is a constant that defines the relative division frequency between these regions. An exhaustive search of the (*f*_*r*_, *y*_*b*_) parameter space was performed to validate the biregional frequency hypothesis.

The orientation of cell division, represented by the angle (*θ*), follows a region-dependent distribution:

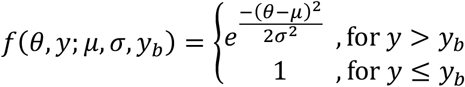

In this framework, *y*_*b*_ is the boundary position. For apical region (*y* > *y*_*b*_), *σ* quantifies the degree of angle bias in division, and *μ* defines the direction of the bias. For the basal region (*y* < *y*_*b*_), the cell division angles were randomly distributed. *μ* was set to π/2 (vertical direction) according to the EdU pulse chase experiment. The specific division angles for cell division were sampled using the acceptance-rejection method. We conducted an exhaustive search of)1/*σ, y*_*b*_) parameter space to test the hypothesis of biregional angles.

For the smooth boundary of biregional biased cell division angle, the function of distribution of cell division angle (*f*(*θ, y*)) was further modified as:

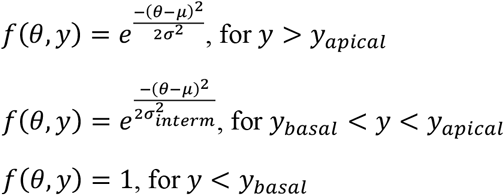

where *y*_*basal*_ and *y*_*apical*_ are the boundary positions of the basal and apical regions, respectively, with an intermediate region between them, and *σ*_*interm*_ is the degree of biased division angles in the intermediate region. 1/*σ*_*interm*_ was set to be a linear interpolation between (1/*σ*_*apical*_, *y*_*apical*_ and (0, *y*_*basal*_), illustrating a gradual shift in biased degree across this intermediate region. 1/*σ* and *μ* was set to be 10 and π/2 under this setting for mimicking complete vertical division. An exhaustive parameter search of (*y*_*apical*_, *y*_*basal*_) was conducted to understand the effects of smooth boundaries and intermediate regions.

### 8. Difference index

The difference index (DI) is used to quantitatively validate the contour similarity between simulated and real organs (*Kinoshita et al., 2022*). This index was calculated by determining the total area enclosed between the outlines of the real and simulated organs. To ensure comparability, we first normalized the sizes of both the real and simulated organs by adjusting their vertical (proximo-distal) lengths to a uniform value of 1. For precise measurements, 101 points were set along both sides (left and right) of each organ’s outline at an even distance of 0.01 intervals in the vertical direction. Then, we measured the horizontal distance (*xd*_*i*_) between the corresponding points on the real and simulated organ outlines. The difference index was computed by summing these horizontal distances and multiplying them by the vertical interval (0.01), resulting in the following formula:

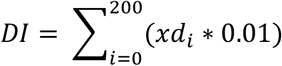

The histogram and heatmap of the difference index of the simulated organ under each specific parameter search and the real organ (mature leaves of *T. sebifera*) were constructed using Gnuplot. Through difference index comparison between real organ shapes (Fig S5), we found that the threshold of 0.10 in difference index shows a high similarity between the simulated and real organ shapes.

### 9. Simulation analysis

All simulations with the same parameters were repeated three times, and the averaged values were used for each analysis. Curvature analysis and difference index calculation used the same algorithm and parameters as described in the experimental data analysis for consistency of the experiment and simulation comparison.

The apical length of the simulated organ was measured as the sudden change in width from the apex to the base. The apex was typically thin and gradually thickened from the apex to the base. When the organ width reached a certain threshold (>30 in the simulation length unit), the apical region was recognized as ending and the apex length was calculated.

### 10. Cell angles and division frequency control in temporal pattern

Arrest front is a widely studied phenomenon that occurs during early leaf development. Previous research quantitatively examined the arrest front in Arabidopsis primordia and found that in the early stage (3DAS∼4DAS), mitotic cells were observed throughout the leaf; in the later stage, the cells in the apical region gradually lost the ability to divide, with only the basal cells dividing (5DAS∼8DAS) (*Kamaza et al., 2010*). In addition, our EdU pulse-chase experiment on *T. sebifera* primordia showed a similar pattern (Fig. 4A-4D). To simulate this phenomenon in the vertex model, the division frequency was controlled using the spatiotemporal function:

In the earliest stage, when all cells within leaf primordia are dividing:

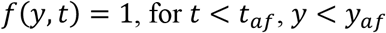

In the next stage, when the arrest front appears, only basal cells can be divided:

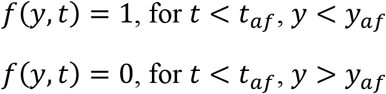

Then in the final stage, when the arrest front shrinks and meristem region decreases:

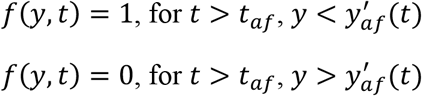

where 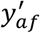 is gradually decreasing over time and is defined by

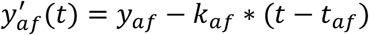

where *k*_*af*_ is the speed of 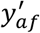 decrease.

Finally, when 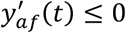, all cell divisions will be arrested and leaf development will transit to the cell expansion stage.

To control the division angle in the temporal pattern, the cell division angles were first set to be completely vertically biased using the Gaussian function:

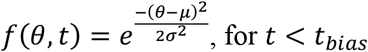

In later stages, the cell division angles were set as random:

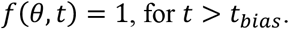

An exhaustive parameter search on (1/*σ, t*_*bias*_) was conducted to test whether temporally biased cell division angles could generate the sharp apex and concave in the simulated organ.

Also, the intermediate pattern was tested for this division angle control:

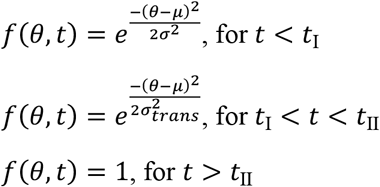

where *t*_I_ and *t*_II_ are the time transition from the vertically biased division angle pattern, random division angle pattern, with an intermediate division angle pattern in between. *σ*_*trans*_ is the degree of biased division angles in the transition. 1/*σ*_*trans*_, was set to be a linear interpolation between (1/*σ, t*_I_) and (0, *t*_II_), illustrating a gradual shift in biased degree across this transition. 1/*σ* and *μ* were set to 10 and π/2 under this setting to mimic vertically biased division during stage I primordia (Fig 4A). An exhaustive parameter search on (*t*_I_, *t*_II_) was conducted to test whether a smooth transition of temporally biased cell division angles could still generate the desired apex and concave shape.

To create a distribution of division angles in temporal control (Fig. 7F, 7G), the position and angle of each division were recorded at every time step and the position was measured as the distance from the tip. At the end of the simulation, the positions were normalized to the final organ shape and the division data were classified into basal and apical regions based on this position. Subsequently, a distribution of angles was created for each region.

## Acknowledgements

We thank Koishikawa and Nikko Botanical Gardens, University of Tokyo, for their support with the sharp-tipped leaf sample collection and *Triadica sebifera* seed collection. We also thank the Ministry of Education, Culture, Sports, Science, and Technology and the Graduate Program for Leaders in Life Innovation (GPLLI)/World-leading Innovative Graduate Study Program for Life Science and Technology (WINGS-LST) of the University of Tokyo for providing the FV3000 confocal microscope facilities.

---

Figure S1 Leaves with characteristic apex. 28 leaves from (from left to right, top to bottom): 1. *Maclura tricuspidate* (Carrière) Bureau, acuminate; 2. *Callicarpa japonica* Thunb., acuminate; 3. *Enkianthus perulatus* (Miq.) C.K.Schneid., mucronate; 4. *Lagerstroemia indica* L., acuminate; 5. *Cornus officinalis* Torr. ex Dur., acuminate; 6. *Vicia unijuga* A.Braun, acute; 7. *Rhododendron kiusianum* Makino, acute; 8. *Ficus erecta* Thunb., mucronate; 9. *Orixa japonica* Thunb., acute; 10. *Rhus succedanea* L., acuminate; 11. *Rhododendron amanoi* Ohwi, mucronate; 12. *Ilex integra* Thunb., acute; 13. *Chimonanthus praecox* (L.) Link, acute; 14. *Lindera communis* Hemsl., acuminate; 15. *Adina pilulifera* (Lam.) Franch. ex Drake, acuminate; 16. *Ruscus aculeatus* L., mucronate; 17. *Cinnamomum camphora* (L.) J.Presl., acuminate; 18. *Distylium racemosum* Siebold & Zucc., acute; 19. *Styrax japonica* Siebold & Zucc., mucronate; 20. *Wisteria sinesis* (Sims) DC., mucronate; 21. *Triadica sebifera* (L.) Small., acuminate; 22. *Impomea coccinea* L., acuminate; 23. *Stephania japonica* (Thunb.) Miers, acute; 24. *Persicaria delibis* (Meisn.) H.Gross ex W.T.Lee, acuminate; 25. *Hibiscus hamabo* Siebold & Zucc., mucronate; 26. *Acer buergerianum* Miq., acute; 27. *Liquidambar styraciflua* L., acuminate; and 28. *Acer palmatum* Thunb., acuminate; The x-axis corresponds to the clockwise directed position of leaf contours and the y-axis corresponds to the curvature. Scale bar 2 cm.

Figure S2. Cell morphology in the mature leaf of *Triadica sebifera*. (A) Mature leaf of *T. sebifera*. (B) six regions of this mature leaf were cut for cell morphology examination. Regions 1 and 2 were defined as the apical region; Regions 3-6 were defined as the basal region. (C) Fluorescence image of epidermal cells under confocal microscopy observation. Cell walls were stained by Calcofluor white. (D) Cell morphology extracted from (C). (D-F, M-O) Cell morphology in regions 1-6. (G-I, P-R) Distribution of cell expansion in (D-F, M-O). (J-L, S-U) The corresponding histogram of (G-I, P-R). Number of cells: (D) 145 cells, (E) 104 cells, (F) 107 cells, (M) 108 cells, (N) 109 cells, and (O) 139 cells.

Figure S3. Cell morphology in the young leaf of *T. sebifera*. (A) The fluorescence image of the apex of a young leaf. Scale bar 50 µm. (B) The distribution of epidermal cell expansion in (A). (C) Distribution of epidermal cell expansion in basal region. (D, E) Corresponding histogram of (B, C). (F) Distribution of epidermal cell expansion in the joint region. (G) Distribution of palisade cell expansion in the basal region. (H) Distribution of palisade cell expansion in the joint region. (I-K) Corresponding histogram of (F-H). Number of cells: (B, D) 113 cells, (C, E) 737 cells, (F, I) 473 cells, (G, J) 723 cells, and (H, K) 666 cells.

Figure S4 Panel of simulation results in biregional biased division angles setting. The right middle part of the panel showed the *T. sebifera* like sharp-apexed shapes. The degree of biased cell division angles in the apical region was defined by the 1/*σ* in the Gaussian function.

Figure S5. The averaged contour and difference index analysis for mature *T. sebifera* leaves. (A) Ten mature leaves of *T. sebifera*. (B) Corresponding curvature pattern of mature leaves. (C) The averaged contour of mature leaves. (D) The corresponding curvature of (C). (E) The difference index between ten mature leaves and the averaged leaves.

Figure S6. Panel of simulation results under biregional biased division frequency setting. All parameters in the exhaustive search failed to generate the desired sharp-apexed shapes (Fig. 5). Some parameters in the upper right part and lower left part generated the Arabidopsis petal-like obovate shapes, which correspond to our previous meristem position simulation research (*Kinoshita et al., 2022*).

Figure S7. Panel of simulation results under biregional division angles setting with gradual boundary. The gradual boundary setting generated a smooth apex.

Figure S8. Temporal control of division angles. (A) Panel of simulation results by the arrest front and temporal division angles combined setting with abrupt angles change (*y*_*arrest*_ = *t*_I_ = *t*_II_). (B) Heatmap of minimum curvature of (A). (C) Heatmap of difference index of (A). (D) Panel of simulation results by the arrest front and temporal division angles combined setting with gradual angles change (*y*_*arrest*_ fixed to be 20).

Figure S9. Cell morphology in mature leaves of *Persicaria debilis* and *Ipomoea coccinea* showed no biased expansion. (A) Mature leaf of *Persicaria debilis*. (A’) Cut region of (A) for cell morphology experiment. (1) is recognized as the apex and (2)–(8) as the base. (B) Cell morphology of apex. (C) Cell morphology of the base. (D) Mature leaf of *Ipomoea coccinea*. (D’) Cut region of (D) for cell morphology experiment. (E) Cell morphology of apex in (D). (F) Cell morphology of base in (D). Scale bar: (A, A’, D, and D’) 1 cm. (B, C, E, and F) 100 µm. The guard cells, trichomes, and their neighboring cells were not included in the expansion analysis because they possess distinct shapes. (B’) Statistical analysis of biased cell expansion of *Persicaria debilis* in the apical region; (C’) Statistical analysis of biased cell expansion of *Persicaria debilis* in the basal region; (E’) Statistical analysis of biased cell expansion of *Ipomoea coccinea* in the apical region; (F’) Statistical analysis of biased cell expansion of *Ipomoea coccinea* in the basal region. Number of cells: (B’) 139 cells; (C’) 318 cells; (E’) 414 cells; and (F’) 695 cells.

Figure S10. Leaves with concave apex and their curvature pattern. (A) *Oxalis lobata* Sims, (B) *Oxalis debilis* Kunth, (C) *Oxalis barsiliensis* Larrañaga, (D) *Oxalis triangularis* A.St.-Hil, and (E) *Liriodendron tuplifera* L. (A’-D’) The curvature pattern of leaflets of the corresponding *Oxalis*. (E’) Curvature pattern of (E). All curvature patterns showed a negative curvature at the apex (at position 50). (F-H) Contour growth mapping results corresponding to (A-C) by controlling the size of non-meristematic regions in near midvein (F: *θ* = 80°, *β* = 0.50, *k* = 0.2; G: *θ* = 20, *β* = 0. 70, *k* = 0.6; H: *θ* = 5, *β* = 0.95, *k* = 1.5). The wider and deeper non-meristematic regions generate wider and deeper concave regions. (F’-H’) Curvature patterns of (F-H). Scale bar: 1 cm.

## Supplementary Movies S1-S16

Supplementary Movie S1: Contour growth mapping of the homogeneous growth settings corresponding to Fig. 2E.

Supplementary Movie S2: Contour growth mapping of vertical growth setting (*β* = 0.8), corresponding to Fig. 2F.

Supplementary Movie S3: Contour growth mapping of biregion growth setting (*β*_*x*_ = 0.60, *β*_*Y*_ = 1.00, *y*_*a*_ = 0.90 *y*_*b*_ = 0.5), corresponding to Fig. 2G.

Supplementary Movie S4: Contour growth mapping of biregion growth setting with fine-tuning (*β*_*x*_ = 0.8, *β*_*y*_ = 1.10, *y*_*a*_ = 1.00, *y*_*b*_ = 0.90), corresponding to Fig. 2H.

Supplementary Movie S5: Vertex model simulation with homogeneous growth setting (biregion division angles, *y*_*b*_ = 0.50, 1/*σ* = 0.10) generated an organ with a circular shape, corresponding to Fig. 4I.

Supplementary Movie S6: Vertex model simulation with vertical growth setting (biregion division angles, *y*_*b*_ = 0.0, 1/*σ* = 10) generated an organ with a rod shape, corresponding to Fig. 4M.

Supplementary Movie S7: Vertex model simulation with biregion growth setting (biregion division angles, *y*_*b*_ = 0. 70, 1/*σ* = 10) generated an organ with a short tip and rounded base, corresponding to Fig. 4J.

Supplementary Movie S8: Vertex model simulation with biregion growth setting (biregion division angles, *y*_*b*_ = 0.60, 1/*σ* = 10) generated an organ with a sharp apex, rounded base, and high similarity to real *Triadica sebifera* leaf, corresponding to Fig. 4K.

Supplementary Movie S9: Vertex model simulation with biregion growth setting (biregion division angles, *y*_*b*_ = 0.30, 1/*σ* = 10) generated an organ with a long tip and rounded base, corresponding to Fig. 4L.

Supplementary Movie S10: Vertex model simulation with a biregion frequency setting (*y*_*b*_ = 0. 7, *rf* = 0.1) generated an organ similar to Arabidopsis petals, as shown in Fig. 5B.

Supplementary Movie S11: Vertex model simulation with the arrest front setting generated an organ with a cylindrical shape, corresponding to Fig. 7C.

Supplementary Movie S12: Vertex model simulation with the temporal division angle setting generated an organ with a rod shape.

Supplementary Movie S13: Vertex model simulation with arrest front and temporal division angle settings generated an organ with a sharp apex and rounded base, corresponding to Fig. 7E.

Supplementary Movie S14: Contour growth mapping of concave growth setting (*θ* = 80°, *β* = 0.50, *k* = 0.2) generated an organ with a wide concave apex, corresponding to *O. lobata* and Fig. S10F.

Supplementary Movie S15: Contour growth mapping of concave growth setting (*θ* = 20, *β* = 0. 70, *k* = 0.6) generated an organ with a narrow and deep concave apex, corresponding to *O. debilis* and Fig. S10G.

Supplementary Movie S16: Contour growth mapping of concave growth setting (*θ* = 5, *β* = 0.95, *k* = 1.5) generated an organ with a narrow and shallow apex, corresponding to *O. barsiliensis* and Fig. S10H.

